# A model predictive control strategy to regulate movements and interactions

**DOI:** 10.1101/2022.08.24.505193

**Authors:** A. Takagi, H. Gomi, E. Burdet, Y. Koike

**Author notes:** **Materials & Correspondence** Correspondence and requests for material should be sent to A.T.

## Abstract

Humans are adept at moving the arm to interact with objects and surfaces. The brain is thought to regulate motion and interactions using two different controllers, one specialized for movements and the other for force regulation. However, it remains unclear whether different control mechanisms are necessary. Here we show that the brain can employ a single high-level control strategy for both movement and interaction control. The Model Predictive Control (MPC) strategy introduced in this paper uses an internal model of the environment to plan the arm’s muscle activity whilst updating its predictions using periodic feedback. Computer simulations demonstrate MPC’s ability to produce human-like movements and after-effects in free and force field environments. It can simultaneously regulate both force and stiffness during interactions, and can accomplish motor tasks demanding transitions between motion and interaction control. Model Predictive Control promises to be an important tool to test ideas of motor control as it can handle nonlinear dynamics with changing environments and goals without having to specify the movement duration.

## Introduction

Humans use their hands to touch and explore their surroundings from as early as the gestation period inside of the mother’s womb [1]. The ability to voluntarily control the arm’s motion, including the force it applies to the outside world, is critical to interacting with the objects that surround us [2,3]. Such interactions are vital to learning from others [4,5], yet the literature on interaction control [6–10] is sparse when compared with free movements that have been studied for over a century. A major difference between free movements and interaction control is in the latter’s unknown dynamics. Interactions can be nonlinear due to friction, and they can be unstable since contact can be easily lost. Furthermore, the elasticity of the surface can vary widely from a soft human hand to a hard smartphone screen. The brain has to control the arm during motion and during contact, and it has to handle transitions between the two. Successful interaction control is difficult to achieve, and it is an area that continues to receive attention from researchers in robotics [11]. Understanding the control strategy utilized by the brain to achieve such versatility during free motion and during contact would have important applications.

Some of the earliest models of human interaction control were inspired by direct force control, which assumes that the arm generates a forward force equal to the desired interaction force [9,12]. However, this idea has fallen out of favor since the arm cannot generate a force like an electric motor with negligible viscoelasticity. Indirect force control has been proposed as a suitable alternative where the arm’s elastic muscles generate the forward force [13,14]. The advantage of this mechanism is that the elasticity of the contact need not be estimated. However, due to its reliance on force feedback signals, continuous contact is an absolute must to guarantee safe and successful operation. Loss of contact can lead to unstable behavior.

To address this limitation of indirect force control, some studies have proposed a blend of force control combined with position control to regulate interactions, possibly switching between the two depending on the presence of contact [15–17]. Force control is primarily used to regulate the interaction force while position control is employed to move the arm without contact. In position control, the brain leverages the spring-like properties of the muscles to pull the arm towards an equilibrium posture. It is proposed that this equilibrium-point is moved to manipulate the arm’s position and reach distant targets. However, switching between control mechanisms makes actions unsmooth [18]. This is especially evident during rapid transitions from free motion to contact, like when using a jackhammer to break concrete, which would require the brain to switch the arm’s control mechanism every time the jackhammer loses contact with the surface. Furthermore, these low-level mechanisms lack the ability to predict environmental dynamics, which is known to be highly effective in achieving movement goals [19].

Here, we propose that the brain does not switch between two different control mechanisms, but uses a single high-level cortical control strategy based on Model Predictive Control (MPC) to regulate the arm during both free motion and contact. The brain may do so by using an internal model of the environmental dynamics to plan the necessary muscle activity ahead of time. The environmental dynamics can be learned in a number of ways using statistical learning techniques such as deep or reinforcement learning [20]. In this paper, we kept the learning process as simple as possible to focus on the arm’s behavior if the brain relies on a predictive model to control it. Comparisons are made between simulations and empirical data to quantify the human-likeness of MPC. The explanative power of the MPC strategy is presented in simulations of two experiments from the literature and an original experiment that tested a prediction of MPC when interacting with a soft environment. These experiments demonstrate MPC’s ability to: (i) move the arm in novel environments, (ii) concurrently control the arm’s elasticity and force, and (iii) control ballistic transitions from free motion to contact. By using a prediction of the environmental dynamics, the brain can control the arm in various environments. MPC receives sensory feedback and updates its motion plan every 150 ms to reflect the central nervous system’s movement generation process which is believed to occur at 7-10 Hz [21,22].

## Results

Figure 1 shows a sketch of the MPC strategy. It finds the muscle activity **u** to achieve the task goal within a receding horizon *T* using minimal effort, which is quantified by the scalar cost function

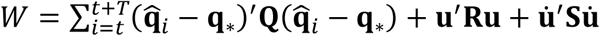

where **Q, R** and **S** are cost matrices, **q**_∗_ is the desired joint state and 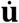 is the change in muscle activity **u**. Because this motion plan is yet to be executed, the future joint state 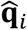 at time step *i* is not measurable, and so it is predicted from the last known state 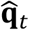 incrementally using 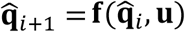 where **f** is a model incorporating the arm and the environment’s dynamics. By doing this for the duration of the receding horizon, the sequence of predicted states 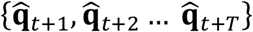 can be obtained. The first of the cost function punishes deviations from the desired joint state, the second minimizes superfluous muscle activity, and the third reduces rapid changes in muscle activity as it demands more energy [23]. MPC solves this Quadratic Programming (QP) problem iteratively to find the muscle activity **u**, which is kept constant over the receding horizon to reduce optimization time. Every *T* seconds, MPC determines a new muscle activity **u** to minimize the cost, thereby fulfilling the task in an unspecified duration.

**Figure 1.**
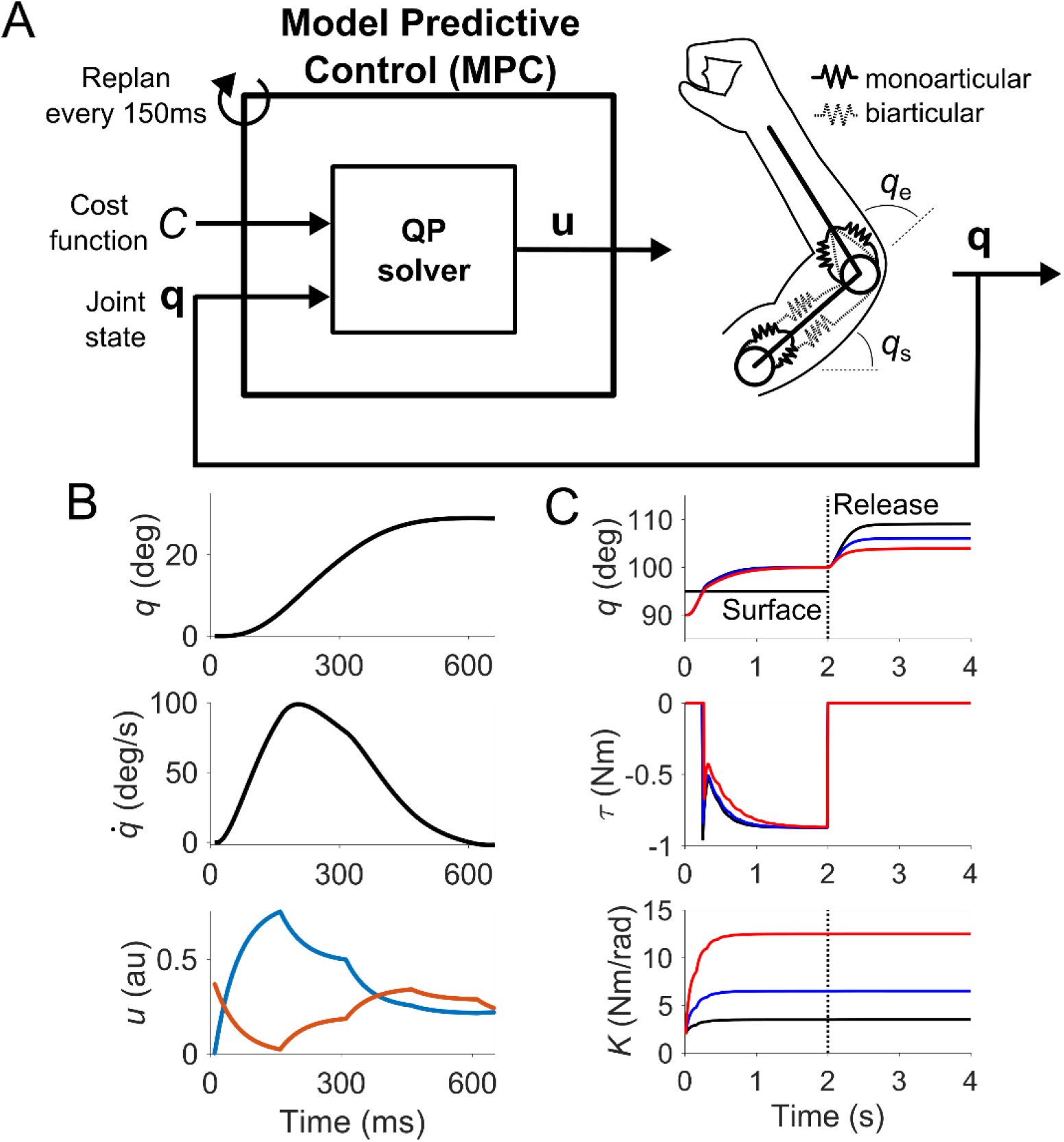
Model Predictive Control uses a model of the arm and the environment to plan the muscle activity required to accomplish a given task. (A) Schematic of MPC finding the muscle activity that minimizes a cost function that defines the goal of the task, e.g. to exert a desired force or reach a desired position. The planning process is repeated once the receding horizon elapses, and continues until the task is fulfilled. The muscle activity is constant for the duration of the plan. (B) Example of MPC moving a single joint through tri-phasic flexion and extension. (C) MPC controlling the single joint system to penetrate a surface positioned at 95°. After five seconds, the surface disappears, causing the joint to move. MPC predicts the joint’s displacement to be shorter when the joint stiffness is stiffer (red) than when it is compliant (blue and black), even when the same force was exerted against the surface prior to its release.

We use a simple musculoskeletal model of the arm for the ensuing computer simulations. Let us first run through an example of MPC controlling a single-joint system by activating a flexor-extensor muscle pair towards a given goal (Figure 1B). The muscle’s elasticity was programmed to increase linearly with its activation [24,25], and the flexor muscle could pull maximally to 180° and the extensor muscle to 0°. The movement range of the joint was between 0° and 180°. Importantly, the torque exerted by each muscle is a function of the joint’s angle and velocity, meaning the torque can change even if the muscle activity is constant. We assumed equal viscoelastic parameters in the flexor and extensor muscles so that the equilibrium posture at null muscle activity was at 90°. To reach the target position, MPC generated a tri-phasic burst in muscle activity [26] through a bell-shaped velocity profile [27]. So MPC can successfully move the joint in a free environment in a human-like manner, but how about during interactions?

Interaction control requires the simultaneous control of the arm’s force and its stiffness, the latter playing a critically important role to stabilize the arm when pushing against a surface [28]. MPC was given the goal to exert a constant force against a surface while simultaneously maintaining a target joint stiffness (Figure 1C). Three conditions were tested where MPC had to keep the joint stiffness low, medium or high whilst exerting the force, which was identical in all conditions. A stiffness goal was explicitly added to the cost function, forcing MPC to coactivate the flexor and extensor muscles whilst exerting the target force. MPC successfully pushed against the surface to reach the target torque (Figure 1C, middle panel) while keeping the joint stiffness at the desired level (bottom panel). If we observed solely the arm’s position and its force after 2 seconds (when their values have stabilized), we cannot spot any difference in the arm’s control despite the arm’s stiffness being different in all three conditions. However, MPC predicts that the arm behavior will be different when the surface abruptly disappears. When the surface is removed at the 2 second mark, the most compliant arm (black) moves and stops far from the original constraint. Compared with the stiff arm (red), the arm with low stiffness travels farther. The arm controlled by MPC is expected to travel a shorter distance from the constraint when the arm’s stiffness is greater when pushing against a surface that suddenly disappears.

An experiment was conducted to test this prediction that the force and the stiffness can be controlled separately by the brain (Figure 2A). We recently showed that participants who increased their grasp strength also increased the coactivation of the shoulder and elbow muscles, causing the arm’s stiffness magnitude to increase linearly with the grasp strength [29]. Using this linear relationship between the grasp strength and the arm’s stiffness, we asked participants to grasp a handle by {4, 7, 10, 13} N. While doing so, they also had to apply a constant 0.7 N force against the handle in one of eight directions (Figure 2B). The position of the handle was kept fixed by powerful motors. Once the desired force and grasp strength were held for 500 ms, the motors were switched off, causing the hand to move like in the simulation of Figure 1C. The hand traced a distinctive path that was dependent on the direction of the force, and the arm became motionless after 800 ms (Figure 2C, black). No voluntary movements were observed after the constraint disappeared.

**Figure 2.**
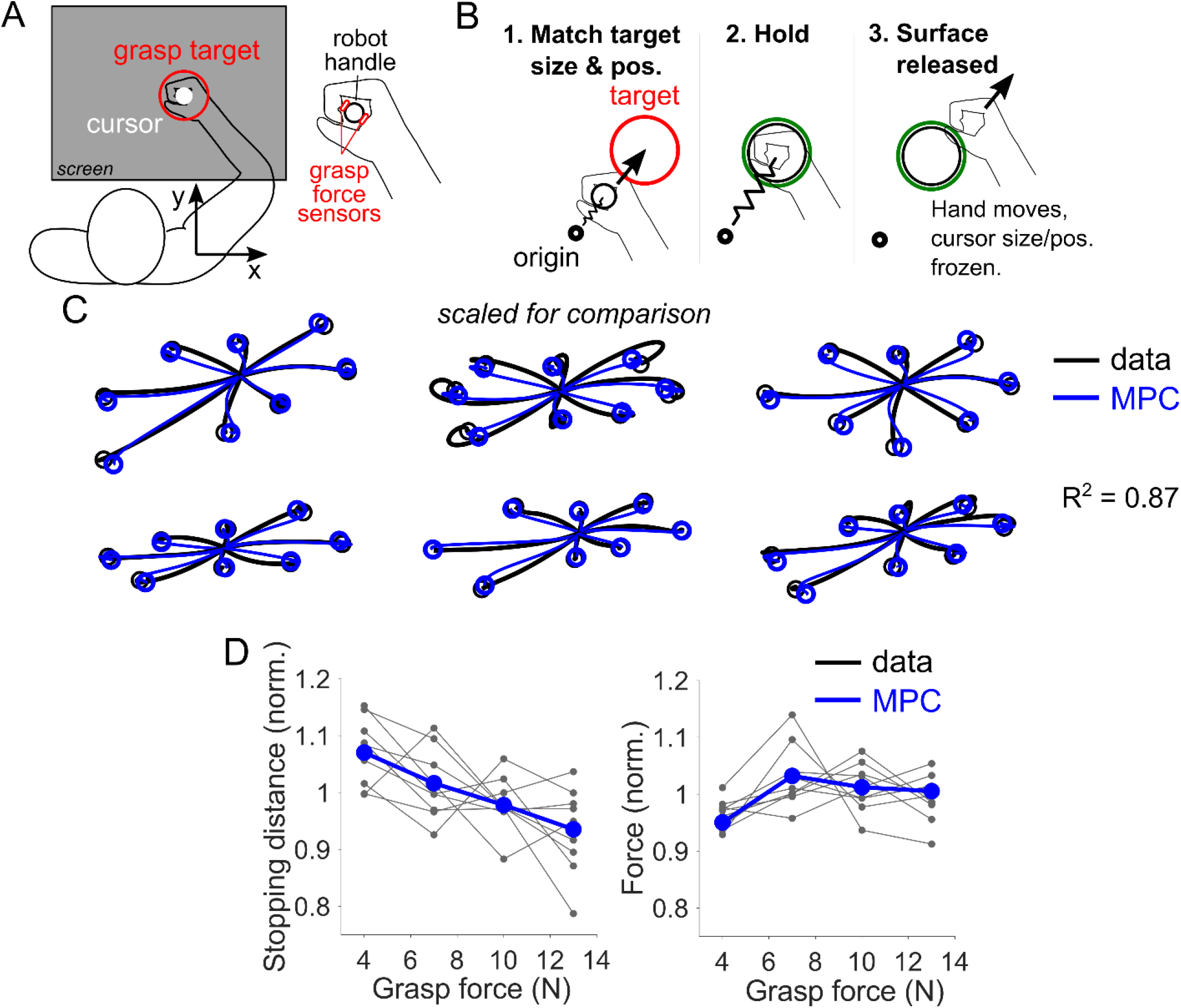
The MPC mechanism accurately reproduces the arm’s trajectory when contact is lost unexpectedly during constant force regulation. (A) Schematic of experimental setup. Participants sat in front of a robotic interface while their grasp force was measured by two three-axis load cells. (B) Participants had to match the position of their cursor (white circle) with the target (red circle) by pushing against a surface and match the cursor to the target size by grasping the handle. After a randomized duration period, the surface disappeared. (C) Predicted and empirical trajectories from six representative participants averaged across all grasp levels. The hand’s trajectory, including its distinct curvature, was accurately predicted by MPC (R^2^ = 0.87±0.02). (D) The normalized force magnitude and stopping distance as a function of the grasp level (black lines are individual data). In line with MPC’s prediction, the hand was displaced by a shorter distance when the arm exerting the same force was stiffer.

We tested whether MPC could accurately predict the hand’s trajectory given the arm’s inertia, viscoelasticity and the force it was applying against the constraint. Since no voluntary movements were discernible once the arm was released, the ensuing motion likely came from the arm’s musculoskeletal dynamics, so each participant’s arm impedance was estimated by linearly regressing the force and position time-series data within an 800 ms time window after the constraint’s disappearance. These values were converted from Cartesian to joint space based on each participant’s posture. The parameters were fed to a simulation wherein MPC found the muscle activity to replicate the arm’s force and stiffness prior to the constraint’s disappearance. MPC had no knowledge of the arm’s trajectory nor the hand’s final position when it came to rest. Nevertheless, it accurately reproduced the arm’s trajectory after the constraint disappeared (coefficient of determination R^2^ = 0.87 ± 0.02, mean±SEM). The distance travelled by the arm or the stopping distance, which was normalized for between-subject comparison, depended on the grasp strength. A one-way repeated measures ANOVA (RM ANOVA) revealed a significant effect of the grasp strength on the stopping distance (*F*(3,24)=5.9, *p*=0.004). The stopping distance was significantly shorter when the grasp force was 13 N compared with 4 N (*p*=0.04, Tukey’s HSD). A stiffer arm stopped closer to the constraint compared with a compliant arm. This cannot be explained by a difference in the applied force, which was constant as a function of the grasp strength (one-way RM ANOVA, *F*(3,24)=1.5, *p*=0.25). Hence the brain can control the arm’s stiffness and force independently of each other in a manner consistent with what MPC expects. However other strategies like equilibrium-point control and impedance control also expect a stiffer arm to come to a stop closer to the constraint. What sets MPC apart from these control strategies?

The MPC strategy differs from equilibrium-point and impedance control concerning the control of movements in free and force field environments. To illustrate our point, we simulated a well-known experiment wherein participants learned to reach eight different targets in a clockwise velocity-dependent force field (VF) [30]. In the original experiment, participants made point-to-point reaching movements to eight targets by holding a robotic manipulandum with the right hand. In the null-field condition where no force was applied to the hand, and participants reached the target in an almost straight line. In the VF condition, the manipulandum applied a force orthogonal to the movement that was dependent on the hand’s velocity. While the hand’s trajectory was initially curved, it gradually became straight again in the after learning. When the force field was removed, returning to the null-field condition, participants exhibited an after-effect of the learning, the hand curving once again but in the opposite direction to the initial exposure phase.

MPC was used to control the arm in the null-field phase, the initial and final VF phases, and in the after-effect phase. MPC’s parameters, specifically those reflecting MPC’s knowledge of the environment, were set to reproduce the four experimental conditions. The simulation assumed no external force in the null-field and initial VF phases (prior to learning). The force field was identified and assumed to be present in the final VF and after-effect phases. In the simulation of the null-field phase, MPC made approximately straight movements towards the target, but the trajectory was curved in the initial VF phase, just like in the experiment (Figure 3A). Straight reaching movements were recovered once MPC had knowledge of the force field in the final VF phase. When the force field was suddenly removed in the after-effect phase unbeknownst to MPC, the arm exhibited an after-effect, curving in the direction opposite to the force field. This after-effect also resembles what was observed in the first trial of the after-effect phase in the original experiment. These simulations suggest that MPC closely resembles the way in which the brain controls the arm during movement when the force field is unknown (initial VF phase) or when it is removed unexpectedly (after-effect phase). Unlike MPC, neither equilibrium-point nor impedance control predict the dynamics of the environment. Thus, they cannot reproduce the after-effects observed after motor learning unless they are explicitly modified to do so. Hence the ability to predict the environmental dynamics and exhibit after-effects is a key difference that sets MPC apart from equilibrium-point and impedance control.

**Figure 3.**
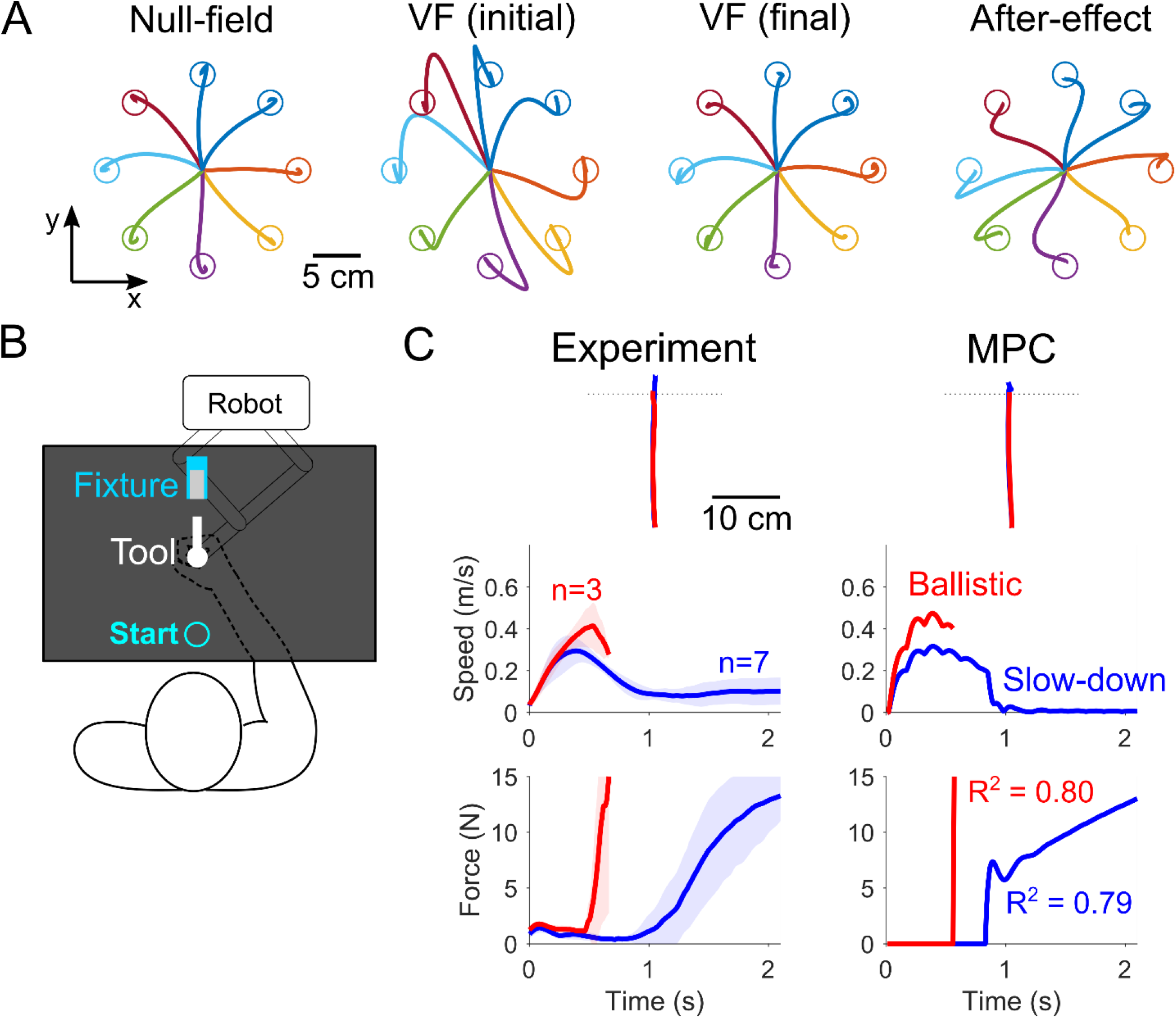
MPC exhibits human-like control during reaching and in tool insertion. (A) Simulation of MPC moving the arm to reach eight targets either in a free environment (null-field and after-effect) or in a velocity-dependent force field (VF). MPC’s trajectories resembles those from the original experiment. (B) Schematic of the tool insertion setup. Participants moved the tool (free motion) and inserted it into the fixture requiring 15 N force. (C) Comparison of the trajectory, speed and force time-series between the empirical data and the best-fitting MPC simulation (see Methods). Since participants in the experiment exhibited two insertion strategies (ballistic and slow-down), MPC was used to predict both strategies, doing so with high accuracy.

So MPC can evidently control the arm during interactions and movements, but can it handle transitions between these two domains? A task like inserting a tool into a fixture, which is a common task in industry, demands such a transition. First, the tool must be transported in front of the fixture, and then it must make contact with the fixture and be forcefully inserted (Figure 3B). In a recent experiment, we demonstrated that participants learning an insertion task adopt one of two strategies [31]. Some participants use a ballistic strategy wherein the tool’s speed barely slowed down as it came into contact with the fixture. The other slow-down strategy was characterized by a slow increase in the force as the tool made contact with the fixture. These two strategies had remarkably different task completion times, the slow-down strategy taking nearly twice as long as the ballistic one, and they could be easily discriminated by their force time-series. Can MPC generate insertion strategies that resemble human-like insertion?

We used the same virtual environment as in the original experiment to assess MPC’s ability to replicate the two insertion strategies (Figure 3B). The two insertion strategies were replicated by different values in the cost function that punished large velocity values. The slow-down strategy had a higher velocity cost than the ballistic strategy, while all other parameters were the same. Monte Carlo simulations were conducted to find the velocity cost parameter that yielded the best fitting force time-series on the population mean data from the ballistic and slow-down strategies separately. The force time-series was accurately predicted by MPC (R^2^ = 0.80 for ballistic, R^2^ = 0.79 for slow-down), reproducing the rapid rise in the force against the fixture during ballistic insertion, and the gradual increase in the force with the slow-down strategy (Figure 3C). MPC also reproduced gross features of the tool’s trajectory in space and its speed time-series. It inserted the tool with a greater peak speed during ballistic insertion, while it used motion with a longer-tailed speed profile in the slow-down strategy, just like in the experiment.

## Discussion

This paper introduced Model Predictive Control as a plausible high-level control strategy for the brain to voluntarily control the arm in diverse environments. Three experiments were simulated that demonstrated human-like control with MPC for (i) interaction control, (ii) movements in free and force field environments, and (iii) facilitation of smooth and rapid transition from free motion to interaction. MPC accurately reproduced or predicted the arm’s behavior in all cases, demonstrating that this single control strategy is capable of handling motion, interactions, and transitions between these domains, and does so with characteristics of the natural motion behavior observed in humans.

MPC replans the muscle activity needed to complete a task every 150 milliseconds, resembling a computational modelling framework recently proposed for biological control systems [32], namely the use of a receding horizon with goals that are updated at a fixed rate [22,33,34]. The replanning of muscle activity at the rate of 7-10 Hz is consistent with submovements observed during the execution of slow movements [21,22], suggesting that the brain may be planning and executing movements at this rate. Different update rates change the size of the after-effects observed during force field learning, with faster updates leading to smaller after-effects. However, the overall control does not change significantly even when the update rate is changed as long as the costs are scaled appropriately to reflect the change in the receding horizon. Unlike other computational approaches like iLQG [35] that optimize and execute for the entire task duration, replanning enables the brain to execute movements without having to specify the task duration. It also helps the brain correct movements on-the-fly in response to sudden changes in the environment or the task, e.g. when the target jumps to a different location [36]. The ability to correct and optimize movements on the fly is particularly important in environments where the dynamics are difficult to predict, which is usually the case when pushing or interacting with surfaces.

Equilibrium-point control and impedance control have also been proposed as plausible mechanisms to regulate both interactions and movements. However, neither mechanism is able to reproduce the after-effects of force field learning unless they are explicitly modified to do so. The after-effect exhibited by MPC is a consequence of it expecting to move in a force field, when in fact it has been removed without its knowledge. The change in the arm’s trajectory during force field learning is a direct consequence of MPC attempting to generate more accurate predictions of its motion plan. This view is consistent with the theory of internal models [37], possibly located in the cerebellum [38,39], and with the recent idea that the brain wants to minimize errors in its prediction or ‘surprise’ [19,40]. In essence, equilibrium-point and impedance control are low-level control mechanisms that do not make predictions about the environment, while MPC is a high-level control strategy that does. Another difference between MPC and equilibrium-point control is in dealing with the problem of muscle redundancy. In the latter, the force against a surface can be increased either by moving the equilibrium-point deeper into the surface or by increasing the arm’s stiffness. However, the mechanism cannot decide which action to take. MPC is different because it takes an energy optimization approach of producing the target force with minimal effort. Thus, it will take the option of least stiffness as this requires least effort.

Studies proposing a hybrid or parallel combination of combination of indirect force control and position control have used differences in brain activity or muscle activity to justify the view that distinct mechanisms are used to control the arm in free motion and interaction [41,42]. However, as muscle activity has to correspond to the environmental dynamics, different neural and muscle activities are expected between arm motion in a free environment and pushing against a surface. Therefore, detecting a difference between these two cannot be used as evidence of distinct control mechanisms. Another study examined interaction control when pushing against a surface that suddenly became soft or rigid [9]. While the authors proposed a parallel force-position controller to interpret the observed behavior, MPC can explain their results. If MPC assumes that the environment is stiffer than it actually is, it will not exert sufficient force. If it believes the surface is more compliant than it is, then it will apply too much force.

The experiment conducted in this study examined how the arm’s behavior changes when it exerts a constant force against a constraint using low or high joint stiffness. By suddenly removing the constraint, we could observe the position to which the hand converged and became stationary, sometimes referred to as its equilibrium posture or attractor. While the equilibrium posture has been estimated in the past [16,43], this is the first instance of it being observed experimentally without relying on extrapolation. The equilibrium posture could be observed due to the small displacement in the arm after the surface disappeared. In preliminary tests, we found that when the arm was displaced by more than 1 cm, like in the study of Kennedy and Schwartz [44], a correctional movement was elicited even though participants were instructed not to do so. The correctional movement elicited a large second peak in the tangential velocity approximately 400 milliseconds after the surface disappeared. This hampered the observation of the equilibrium posture, and why the displacement had to be small during our experiment. A correctional movement could explain why the value of the arm’s impedance estimated in the experiment of [44] is nearly double the reported value from other studies [24,29,45].

There is ample evidence in the literature of the brain learning the dynamics of complex environments, ranging from velocity-dependent force fields [30] to novel objects [2]. Since there exist multiple approaches to estimating and learning the dynamics of a novel environment, a nonlinear Kalman filter being a good example [4], this was kept out of the scope of the current study to focus on MPC’s ability to generate human-like behavior in diverse environments. As the MPC strategy uses a predictive model of the dynamics to plan and execute movements, it naturally fits into the predict, update and correct methodology used in many learning frameworks [46], making it relatively straight-forward to apply a learning principle to MPC. Even the structure of the internal model used to predict the future state of the limb is flexible. In the current study, we opted for a nonlinear parametric model of the arm-environment system, but other models like a neural network can be used in its place. The only requirement is that the model must generate a prediction of the state that can be used to optimize and plan the movement.

The computational approach used in this study is deterministic and not stochastic, and so in its current form, the MPC strategy does not increase coactivation in response to unstable environments, instead choosing to correct for errors using reciprocal activation of the muscles. However, stochastic MPC, which minimizes a cost function composed of the state’s mean and its covariance [47], is capable of increasing the joint stiffness to counter instability. MPC optimizing for a stochastic environment will embody the characteristics of human-like control as shown in this study, and will exhibit human-like stabilization of the arm in unstable environments [48]. Even though the ability to handle unstable environments was out of the scope of this study, stochastic MPC would increase muscle coactivation in the presence of unstable force in a human-like manner according to the simulation study of [48].

MPC is a high-level voluntary control strategy that yields human-like motion planning in free and force field environments, and during interactions with surfaces and constraints. At its core, it yields a motion plan identical to optimal control, e.g. linear quadratic regulators for an infinite replanning time and linear dynamics, and has a structure familiar to computational theories in motor control. MPC is powerful because it does not linearize or approximate the nonlinear dynamics as iLQG does [35], and is significantly more robust in handling constraints to ensure that muscle activity is never negative. Model Predictive Control is a promising computational approach to test new theories of motor control.

## Materials and Methods

### Model of the viscoelastic arm

The arm was modelled as a two-joint, six muscle system with the shoulder’s Cartesian position fixed in place. The shoulder and elbow joints each have two monoarticular muscles that pull in opposite directions. In addition, two biarticular muscles exert a torque simultaneously on the shoulder and elbow joints. Each muscle is modelled as a linear spring-damper that connects the joint segment to its corresponding joint limit angle. As an example, for a shoulder with angle *q*_*s*_ and velocity 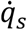, the shoulder flexor with the joint angle limit *q*_*sF*_ = 180° exerts the joint torque

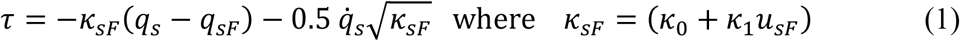

where *κ*_0_ and *κ*_1_ are a stiffness constant and slope, and *u*_*sF*_ ≥ 0 is the muscle activity of the shoulder flexor. Combined with a shoulder extensor with joint angle limit *q*_*sE*_ = 0° and muscle activity *u*_*sE*_ ≥ 0, giving it a stiffness of *κ*_*sE*_ = (*κ*_0_ + *κ*_1_*u*_*sE*_), the total torque on the shoulder is given by

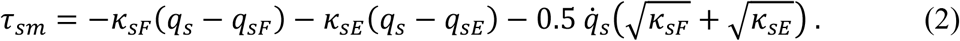

when the muscles are relaxed and if the muscle stiffness parameters are equivalent, the shoulder joint converges to the mean of the joint limits (*q*_*sF*_ + *q*_*sE*_)/2. However, this attractor angle changes depending on the muscle activities *u*_*sF*_ and *u*_*sE*_. It should be noted that the stiffness of the joint can be increased by coactivating the flexor and extensor muscles equally, which does not affect the attractor angle.

The two monoarticular muscles at the elbow with joint limits *q*_*eF*_ = 180° and *q*_*eE*_ = 0° give rise to a joint torque at the elbow in a similar form as eq. (2),

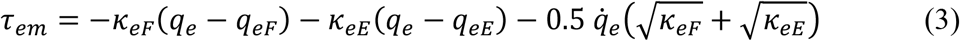

The biarticular muscles exert a torque dependent on both the shoulder and elbow angles *q*_*s*_ and *q*_*e*_ and their respective velocities with joint angle limits identical to the monoarticular muscles. The torque exerted by both the biarticular flexor and extensor muscles is

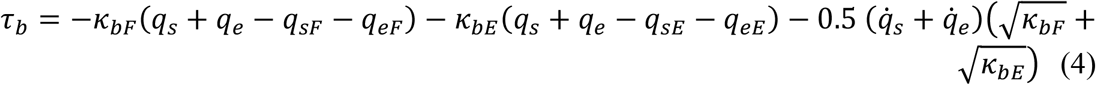

where *κ*_*bF*_ = (*κ*_0_ + *κ*_1_*u*_*bF*_) and *κ*_*bE*_ = (*κ*_0_ + *κ*_1_*u*_*bE*_) are the stiffness of the biarticular flexor and extensor muscles, respectively.

Summing the contributions of eqs. (2), (3) and (4), the six muscles altogether exert a torque **τ** on the arm such that its state **q** = [*q*_*s*_, *q*_*e*_]^′^ evolves in time according to

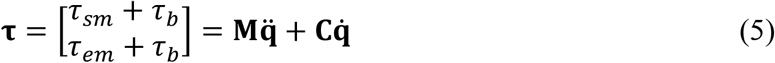

where the arm’s inertia in joint space

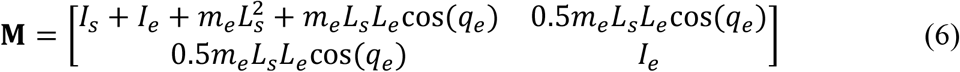

is a function of the elbow angle and

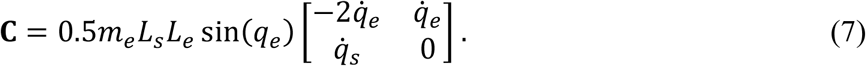

is the Coriolis force. The parameters *L*_*s*_ = 0.3 m and *L*_*e*_ = 0.35 m are the lengths of the shoulder and elbow segments, *I*_*s*_ = 0.05 kgm^2^ and *I*_*e*_ = 0.06 kgm^2^ are their moments of inertia, and *m*_*s*_ = 1.9 kg and *m*_*e*_ = 1.7 kg are their point masses [17]. The joint inertia assumes that the location of the elbow’s centre of mass is halfway along its segment.

### Model Predictive Control

Model Predictive Control uses a model of the arm and the environment to plan the future muscle activity needed to accomplish a given goal. Based on evidence that the brain plans and executes movements at a rate of 7-10 Hz [22,33], MPC plans the muscle activity for a future period of time henceforth referred to as the receding horizon *T*. Since the task does not end once *T* seconds has elapsed, the motion is replanned every *T* seconds until the task is complete. The arm’s super-state 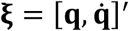 is composed of the two-link arm’s joint positions and velocities, and the goal is to move this arm to a new position and a set velocity **ξ**_∗_. If the current time is *t*, then the cost function *W* takes the form

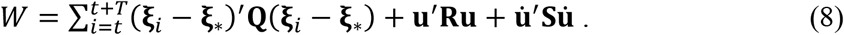

The first term, which is weighted by **Q**, ensures that the joint’s predicted state **ξ**_*i*_ is as close to the desired state **ξ**_∗_. The second and third terms, weighted by **R** and **S** respectively, ensure unnecessary muscle activity and rapid changes in muscle activity are punished to conserve energy [23]. For the current purpose, these weights were the same for all muscles, rendering them as scalar constants *R* and *S*. The muscle activity is constant for the duration of the horizon as this simplifies the optimization process. While the muscle activity may be constant for the horizon, any change in the joint’s position or its velocity will result in a change in joint torque in accordance with eq. (5). Unless specified, the values for the weighting parameters were set to **Q** = diag(200, 200, 1, 1), *R* = 10^−8^, *S* = 1.

MPC predicts the joint state **ξ** iteratively by starting from its last measured value, and iterating it forwards in time using the dynamic model **ξ**_*i*+1_ = **f**(**ξ**_*i*_, **u**). Here, we used a parametric model for the internal model **f**, and the Euler method was used to update the state,

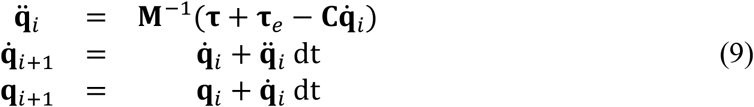

The optimization problem is to find the muscle activity **u** ≥ **0** that minimizes the cost function of eq. (8) using eq. (6). This was solved by Quadratic Programming (QP), namely the iterative updating of the muscle activity until the solution converges to an acceptable minimum. There are several algorithms for solving QP class problems, and the one we opted for was the interior-point method implemented in MATLAB’s ‘fmincon’ function (using its default settings) due to its speed and robustness.

Every instance that *T* seconds elapses, a new motion plan with a new muscle activity **u** is found. The planned muscle activity was filtered by a first-order low pass filter with a time constant of 50 milliseconds to reproduce the filter-like properties of the muscle [49]. All simulations were discretized with a time step of *dt* = 0.0005 s. The arm’s initial posture is specified for each interaction scenario simulation. Non-zero muscle activity is needed to sustain the arm at a given initial posture, and so ‘fmincon’ was used to find the initial muscle activity that produced zero torque at a given initial joint posture.

### Experimental procedure of pushing against a disappearing surface

The procedure of the experiment involving human participants was in accordance with the ethical standards of the institutional research committee (Ethical Review Board for Epidemiological Studies at the Tokyo Institute of Technology, no. 2017087). 9 male participants (25±1 years old), who all gave written informed consent prior to the experiment, participated in the study.

Each participant was seated facing the planar robotic manipulandum (KINARM, BKIN Technologies) with the arm supported by a moving rest underneath the elbow (Figure 2A). The task was conducted with the arm in the horizontal plane at shoulder height. Participants held the KINARM’s handle, equipped with a six-axis force sensor, to exert a force in the two-dimensional workspace. The grasp force was measured using two three-axis force sensors (Tec Gihan) which were positioned at the second phalange of the middle finger and the palm of the hand. The grasp force was defined to be the minimum value of the two sensors. Participants received visual feedback of the task by looking at the reflection of the monitor on a thin-film mirror placed above the robot’s workspace, which obscured the hand and the arm from direct view. They rested their forehead on a frame at a comfortable position and were instructed to keep their head and torso in the same posture throughout the experiment. The torso was constrained by pressing the chest against a fixed table placed below the robotic interface. The position of the shoulder and the lengths of the forearm and the upper arm were measured prior to the task. The position of the robotic manipulandum was controlled to maintain it at the origin of the task [*x*_0_, *y*_0_]′ using a position controller that generated the environmental force 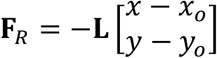. Participants had to push against this constraint to move the cursor on the screen.

A red circle was shown on the display, indicating the target force and target grasp force. Participants had to push against the surface to match the cursor (white circle) and target positions. The position was scaled such that 1N corresponded to 1cm. The cursor grew in size with the grasp force. Once the participant held the force and the grasp force for 700 ms within tolerances of ±0.2 N and ±1 N, respectively, the constraint disappeared after a randomized period between 300 and 800 ms. During this time, the cursor’s position was frozen in place to the last value measured prior to the disappearance, preventing participants from reacting to the sudden movement [50]. The cursor size was updated during this period. Participants could relax 1.5 seconds after the surface disappeared. Participants were instructed to ignore the arm’s movement and focus on maintaining the force and the grasp force. The size of the cursor and the target were rescaled every trial such that the target’s diameter was 1cm in all trials. The target grasp force was one of {4,7,10,13}N. The magnitude of the target force was 0.7 N, and its direction was one of {0 °,45 °,90 °,135 °,180 °,225 °,270 °,315 °}. A target grasp force and target direction pair were drawn randomly every trial, such that each block consisted of 4(grasp levels) x 8(directions) = 32 trials. Each pairing of the target force direction and target grasp force was tested three times in three consecutive blocks, totaling 96 trials. The hand’s velocity, acceleration and the interaction force were low-pass filtered using a zero-lag, second-order Butterworth filter with a 20 Hz cutoff frequency. All data was sampled and recorded at 1000 Hz.

To simulate the empirical results from this experiment, we had to identify each participant’s joint inertia 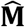, viscosity 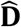 and stiffness 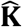. This was done by linearizing the dynamics of the arm in Cartesian coordinates, estimating the arm’s impedance parameters, then converting them to joint space. If the participant pushes against a surface with force **f**_0_ at position **h**_0_ using a grasp force *g*, and then the surface suddenly disappears, the measured force **f** is equal to

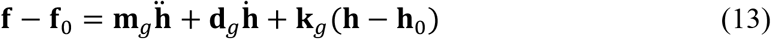

where **h** is the position of the hand in Cartesian coordinates. 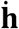 and 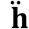 are computed through numerical differentiation, while **f** and **h** are measured within the window between 0 ms and 800 ms after the surface disappears. The data was pooled together from all trials for each participant, and was separated by the four target grasp force levels. Least-squares regression was used to calculate **m**_*g*_, **d**_*g*_ and **k**_*g*_, yielding four values for each participant (for each target grasp level). The amount of variance explained by the fitted parameters was R^2^ = 0.84±0.01, suggesting that the linear model was appropriate in estimating the dynamics of the arm. Only the stiffness magnitude increased with the grasp force, while the inertia and the damping remained constant.

The Cartesian stiffness, viscosity and inertia were transformed to the joint space via the Jacobian **J**

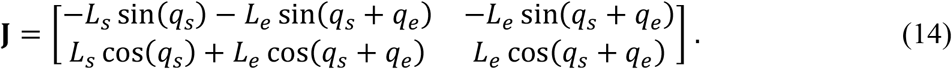

Transforming the arm’s inertia, viscosity and stiffness from Cartesian to joint coordinates yields **M**_*g*_ = **J**^′^**m**_*g*_**J, D**_*g*_ = **J**^′^ **d**_*g*_**J** and **K**_*g*_ = **J**^′^**k**_*g*_**J**. For the last conversion to Cartesian stiffness, zero force was assumed. These parameters were used in the subsequent simulation of the disappearing surface experiment described below.

### Interaction simulation 1: pushing against a disappearing surface

We simulated our experiment where participants pushed against a disappearing surface using a specified grasp force (Figure 2). The arm’s initial posture was set to the value measured from each participant prior to the experiment (*q*_*s*_ = 44 ± 2 degrees and *q*_*e*_ = 85 ± 3 degrees, mean±SEM). Since the task required participants to push against the surface with a target force and a target arm stiffness, the cost function of eq. (8) was modified to

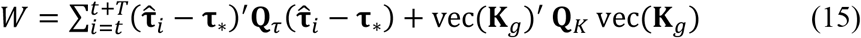

where 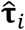 is the torque predicted using the internal model of eq. (9), and **τ**_∗_ is the target torque. The last term **Q**_*K*_ denotes the cost in matching the vectorized joint stiffness vec(**K**_*g*_) estimated from our data. The term punishing muscle activity is omitted because it prevents the muscle coactivation from reaching the target stiffness.

The cost function was dependent on both the participant and the target grasp level, while the force direction was changed by rotating the desired torque **τ**_∗_ to the eight directions of the task. The desired torque is given by transforming the measured force using the Jacobian, **τ**_∗_ = **J**^′^**f**_0_. For all simulations, **Q**_*τ*_ = diag(1, 1) and **Q**_*K*_ = diag(1, 1, 1, 1). The hand’s trajectory from our simulation and the experiment were compared to assess the goodness of MPC’s prediction, which revealed that MPC explained R^2^ = 0.88 ± 0.02 of the variance in the empirical trajectory (Figure 2C).

### Interaction simulation 2: planar reaching in null and force fields

To demonstrate MPC’s human-like control of movements, we simulated the hand’s trajectory during planar reaching in the seminal experiment of [30]. In the experiment, a robotic manipulandum was used to create two virtual environments. In the null field, no force was exerted on the user’s hand, while the velocity-dependent force field (VF) exerted a lateral force to the hand perpendicular to the movement direction. The size and direction of the force **F**_*e*_ depended on the hand’s velocity in Cartesian coordinates,

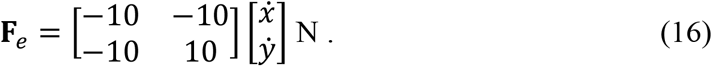

Participants had to reach a target located 10 cm away from their starting position, and had to stop inside the target. The target could appear in one of eight locations spaced equally around a circle encircling the starting position. The experiment was split into four phases: the null-field, the initial VF, final VF, and the after-effect phase. The VF was applied only in the initial and final VF phases. To reproduce their results, MPC’s goal was to minimize the cost function of eq. (8), and its internal model, described by eq. (9), was different for each phase. In both the null-field and the initial VF phases, MPC believed that the environment force was **F**_*e*_ = **0**, and during the final VF and after-effect phases it thought that the external force was equal to the one given in eq. (16). And so in the null-field and final VF phases, MPC’s knowledge and the real environment are the same, but in the initial VF and after-effect phases a mismatch occurs. This gives rise to the trajectories observed in Figure 3A.

### Interaction simulation 3: planar tool insertion

In the original planar insertion experiment, participants used a robotic manipulandum that realized a virtual tool with a length and width of 10 cm and 0.5 cm, respectively, which had to be inserted inside a fixture whose width was 1 cm [31]. The robot generated forces emulating the contact between the tool and the fixture. Outside of the fixture, the tool was in free motion. When the tool’s tip *y* entered the fixture placed at *y*_1_, the robot generated a force

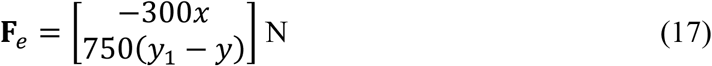

which pushed back against the tool, forcing participants to push against the fixture to insert it fully. The back of the fixture was made of an even stiffer material, which pushed the tool back when it penetrated into the back-wall of the fixture with a force of

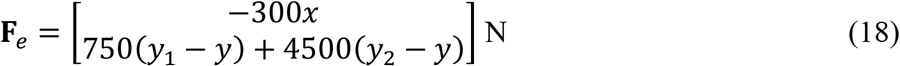

where *y*_2_ is the position of the back-wall of the fixture. MPC’s internal model was set to reflect these values in eq. (9).

The cost function of eq. (8) was minimized by MPC to find the muscle activity to insert the tool into the fixture. To reproduce the ballistic and slow-down insertion strategies, the velocity cost *ω* ∈ [0, 0.04] was incrementally changed within the state cost **Q** = diag(0.3, 0.3, *ω, ω*), which punished differences between the predicted joint state and the desired joint state. The velocity cost parameter that produced a force time-series with the highest R^2^ was selected for Figure 3C. MPC reproduced both insertion strategies, suggesting that the only difference between the ballistic (best value *ω* = 0) and slow-down strategies (best value *ω* = 0.01) was in how fast the tool was allowed to move during the task, the latter being more punishing of large tooling speeds.

## Acknowledgments

The authors would like to thank Domenico Formica, Gowrishankar Ganesh, Neville Hogan, Hiroyuki Kambara and Ted Milner for their comments on an earlier version of the manuscript. All authors conceived the experiment. A.T. collected the data, carried out their analysis, carried out the simulations, and prepared the first version of the manuscript. All authors developed the computational model and revised the manuscript. A.T. gratefully acknowledges funding from the World Research Hub Initiative (WRHI), Institute of Innovative Research (IIR), Tokyo Institute of Technology, Tokyo, Japan. A.T. was partially supported by the JST PRESTO grant JPMJPR18J5. A.T. and Y.K. were partially supported by the JST Mirai JPMJMI18C8 and JSPS KAKENHI 19H05728 grants. E.B. was supported by the European Commission H2020 ICT 871767 REHYB grant.

## Competing interests

The authors have no competing financial or non-financial interests.

## Supplementary Materials

**Supplementary Table 1.**
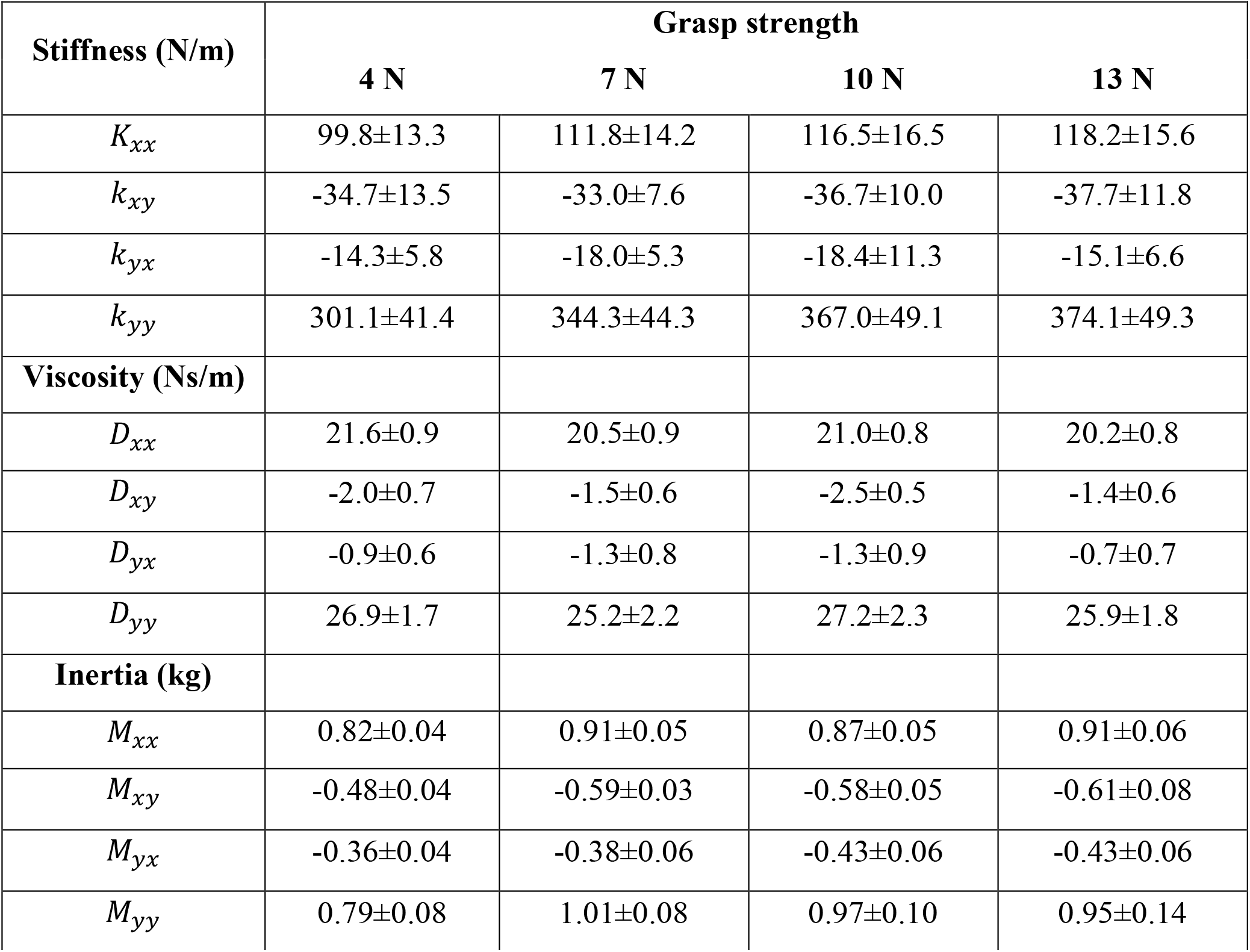
The values of the mean and standard error of the arm’s endpoint stiffness, viscosity and inertia matrix parameters in Cartesian coordinates estimated from the participants in the release experiment. The columns denote the grasp strength. Only the arm’s stiffness was significantly dependent on the grasp strength.

| Stiffness (N/m) | Grasp strength |  |  |  |
| --- | --- | --- | --- | --- |
|  | 4 N | 7 N | 10 N | 13 N |
| $K_{xx}$ | 99.8±13.3 | 111.8±14.2 | 116.5±16.5 | 118.2±15.6 |
| $k_{xy}$ | -34.7±13.5 | -33.0±7.6 | -36.7±10.0 | -37.7±11.8 |
| $k_{yx}$ | -14.3±5.8 | -18.0±5.3 | -18.4±11.3 | -15.1±6.6 |
| $k_{yy}$ | 301.1±41.4 | 344.3±44.3 | 367.0±49.1 | 374.1±49.3 |
| **Viscosity (Ns/m)** |  |  |  |  |
| $D_{xx}$ | 21.6±0.9 | 20.5±0.9 | 21.0±0.8 | 20.2±0.8 |
| $D_{xy}$ | -2.0±0.7 | -1.5±0.6 | -2.5±0.5 | -1.4±0.6 |
| $D_{yx}$ | -0.9±0.6 | -1.3±0.8 | -1.3±0.9 | -0.7±0.7 |
| $D_{yy}$ | 26.9±1.7 | 25.2±2.2 | 27.2±2.3 | 25.9±1.8 |
| **Inertia (kg)** |  |  |  |  |
| $M_{xx}$ | 0.82±0.04 | 0.91±0.05 | 0.87±0.05 | 0.91±0.06 |
| $M_{xy}$ | -0.48±0.04 | -0.59±0.03 | -0.58±0.05 | -0.61±0.08 |
| $M_{yx}$ | -0.36±0.04 | -0.38±0.06 | -0.43±0.06 | -0.43±0.06 |
| $M_{yy}$ | 0.79±0.08 | 1.01±0.08 | 0.97±0.10 | 0.95±0.14 |

## Notes

### Competing Interest Statement

The authors have declared no competing interest.

